# Integrative kinomics reveals kinase-driven subtypes of gastric adenocarcinomas with sunitinib as a potential common inhibitor

**DOI:** 10.1101/2020.02.12.944371

**Authors:** Maria Johanna Tapken, Georg Martin Haag, Dirk Jäger, Stefan Fröhling, Andreas Mock

**Affiliations:** Department of Medical Oncology, National Center for Tumor Diseases (NCT) Heidelberg, Heidelberg University Hospital, Heidelberg, Germany; Department of Translational Medical Oncology, National Center for Tumor Diseases (NCT) Heidelberg, German Cancer Research Center (DKFZ) Heidelberg, Germany; German Cancer Consortium (DKTK), Heidelberg, Germany

**Keywords:** Molecular subtypes, TCGA, precision oncology, targeted therapy

## Abstract

Molecular subtyping of tumors promises a personalized stratification into different treatment regimens. In gastric adenocarcinoma, the four molecularly defined subtypes *chromosomal instable (CIN), microsatellite unstable (MSI), genomically stable (GS)* and *EBV-positive* subtype have been proposed. Following an integrative kinomics approach this computational analysis aimed to predict the best kinase inhibitor for every molecular subtype of gastric adenocarcinomas using publicly available TCGA data (n=404). Intriguingly, using the regulatory network of gastric cancer to estimate protein activity, 43% of all samples could be identified to be kinase-driven. These samples were divided into three clusters with mutually exclusive kinase activities that were independent of the established molecular subtypes. Integrating the pattern of kinase overexpression with an unsupervised target landscape of 37 approved clinical kinase drugs revealed that sunitinib had the best target spectrum across the activated kinases in all three sample clusters. Future work is warranted to validate the kinase-driven subsets of gastric cancer and sunitinib as a potential common inhibitor.

## Introduction

The 2019 WHO classification of gastric adenocarcinomas relies like the Laurén classification from 1965 still on histology. Tubular cell patterns are differentiated from papillary, poorly cohesive (including signet-ring cell and other subtypes), mucinous and mixed adenocarcinomas. While a first molecular classification dividing gastric cancers into four major molecularly defined subtypes has already been proposed in 2014 as part of the TCGA project (Bass et al., 2014), we are still lacking an integrative classification of gastric adenocarcinomas. In contrast to the successful integrative WHO classification of central nervous system tumors (Dewitt, Mock, & Louis, 2017; Louis et al., 2016) this might be due to fact that the gastric cancer subtypes are not defined by single genomic alterations, but by transcriptomic signatures which are yet to be applied in routine pathology workups.

Transcriptomic subtypes however enable a system-wide understanding of oncogenic signaling, as well as the tumor microenvironment paving the way for molecularly guided treatments (Chia & Tan, 2016). A major limitation of RNA-centric precision oncology is that abundant mRNA transcripts do not necessarily translate to a high protein activity. Modern statistical methodologies, including the so-called master regulator analysis, could however show to overcome this limitation by using a network-biology approach to estimate protein activity from transcriptomic information (Alvarez et al., 2018; Califano & Alvarez, 2017).

Kinase inhibitors (KI) as one of the many compounds with anti-cancer effect are sometimes considered dirty drugs because of their broad and often vaguely known inhibition profile. However, recent studies have systematically analyzed binding sites of clinically tested KIs by chemical proteomics. The largest effort till date used a kinobead pulldown in combination with quantitative mass spectrometry to reveal an unsupervised target landscape of clinical kinase drugs (Klaeger et al., 2017).

The main objective of this study was to integrate the oncogenic pathway dependencies in gastric adenocarcinoma based on estimated protein activity using computational inference with the target landscape of clinical kinase drugs.

## Material and Methods

### RNA-seq data analysis of TCGA gastric adenocarcinomas

The Cancer Genome Atlas (TCGA) RNA-seq data (RNASeq2GeneNorm; gene-level data; 19921 genes) of 450 gastric adenocarcinomas (TCGA entity code: STAD – stomach adenocarcinoma) was retrieved through the *curatedTCGA* R package. TPM values were calculated from the count matrix. Updated clinical metadata and the annotation regarding the TCGA immune subtypes (C1-C6) were retrieved from supplementary data of the TCGA pan-cancer initiative (Thorsson et al., 2018). The STAD molecular subtype information was taken from the most recent TCGA study of gastrointestinal cancers (Liu et al., 2018). The molecular subtype was available for 412 of 450 STAD samples. An additional 8 samples belonging to the HM-SNV group were excluded to the small subtype group size.

### Estimation of protein activity using the viper algorithm

The transcriptional regulon of gastric adenocarcinomas was downloaded through the R package *arcane.networks*. The VIPER algorithm was run with the STAD regulon and the viper function from the *viper* R package with default parameters (Alvarez et al., 2016). Using the VIPER algorithm, the protein activity of in total 5275 genes could be estimated (Fig. 1A). A significant protein activity was defined as the top 5% quintile of estimated protein activities (EPAs).

**Fig. 1.**
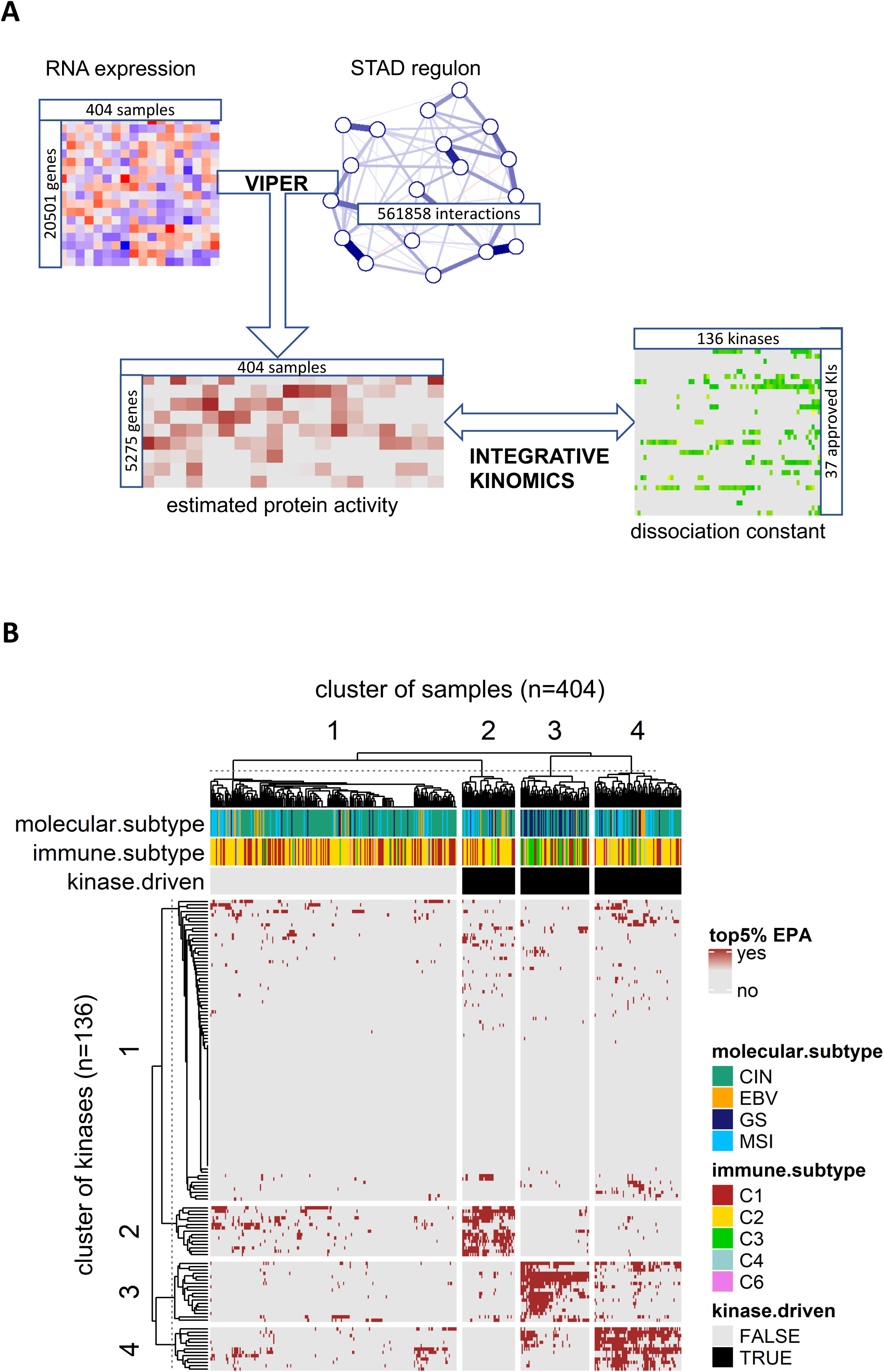
Identification of kinase-driven STADs by integrative kinomics. (A) Graphical abstract of the integrative kinomics approach. First, protein activities were estimated by the so-called viper algorithm that combines RNA-seq tpm values with the regulatory network of gastric adenocarcinomas (i.e. regulon). Next, the protein activity is integrated with the unsupervised target landscape of clinical kinase drugs. (B) Heatmap of protein activity landscape of 136 kinases in the 404 TCGA STAD samples. The column annotation shows the four TCGA molecular subtypes and six TCGA pan-cancer immune subtype (C1-C6). Clusters were defined by k-means clustering with k=4 for both rows and columns. Overactivity of a kinase was defined as belonging to the top 5% of estimated protein activities (EPAs). CIN, chromosomal instability; EBV, Epstein-Bar virus; GS, genome stable; KI, kinase inhibitor; MSI, microsatellite instability; STAD, stomach adenocarcinoma.

### Integrating the target landscape of clinical kinase drugs

The target landscape of clinical kinase drugs was obtained from the work of Klaeger and colleagues (Klaeger et al., 2017). The matrix of apparent dissociation constants (KDapp in nM) was retrieved from the supplemental material of the publication. The 37 approved clinical kinase drugs bound to in total 136 different kinases.

### Statistical analysis and data visualization

Unsupervised analysis was performed by t-distributed stochastic neighbor embedding (t-SNE). Linear modelling (*limma* R package) was used to compare the EPAs between the molecular subgroups. Cutoffs for a significant difference were an FDR-corrected p-value less than 0.01 and a log2 fold change of greater than 0.25. Heatmaps were generated using the *ComplexHeatmap* R package (Gu, Eils, & Schlesner, 2016). Enrichment of sample clusters with molecular or immune subtype was analyzed by Fisher’s Exact Tests.

## Results

### Comparative analysis of the estimated protein activity in gastric adenocarcinoma subgroups

A total of 404 stomach adenocarcinoma samples from the TCGA cohort were analyzed in this study. Among them were 242 classified as chromosomal instable (CIN), 78 microsatellite instable (MSI), 53 genomically stable (GS) and 31 Epstein-Barr virus (EBV)-positive tumors (Fig. S1a). T-distributed stochastic neighbor embedding (t-SNE) performed on the 5000 most variant genes on the RNA level (Fig. S1B), as well as the 5275 genes for which an estimation of protein activity was possible (Fig. S1C) did not suggest a transcriptome-wide difference between the gastric adenocarcinoma molecular subgroups. A supervised comparison of estimated protein activities (EPAs) however revealed subtype-specific overactivity, i.e. 172 genes in CIN, 163 genes in EBV, 105 in GS and 271 in MSI tumors (Fig. S2).

### Kinase activity landscape reveals three subsets of kinase-driven gastric adenocarcinomas

While the molecular subtypes displayed differences in protein activity, the main incentive of this analysis was to investigate the activity of the 136 kinases that are targets for the 37 approved kinase drugs. As a conservative approach, a significant activity was defined as an EPA belonging to the 5% quintile of all EPAs within the cohort. Hierarchical clustering revealed in total four sample clusters that were defined by four different clusters of kinase expressions (Fig. 1B). 43% of samples belonging to clusters 2,3 and 4 did indeed show elevated kinase activity and were hence denoted to be kinase-driven. 57% of tumors on the other hand didn’t display a pattern of kinase overexpression. Cluster 3 contained 63 samples that were defined by activity of 18 kinases, was enriched for GS tumors and an inflammatory (C3) immune subtype (p-value 1.5E-11 and 3.0E-12, respectively; Fisher’s exact test). The remaining clusters did not show a significant association with a STAD molecular or immune subtype.

### Integrative kinomics reveals sunitinib as a potential common inhibitor of kinase-driven tumors

Next, the target landscape of 37 approved kinase inhibitors was interrogated for the three different kinase-driven subsets (Fig. 2; cluster 2 – 15 kinases, cluster 3 – 18 kinases, cluster 4 – 13 kinases). Here, we were most interested in drugs that showed an affinity for as many kinases as possible within a cluster. Intriguingly, while the three clusters harbored mutually exclusive kinase activities, the multikinase inhibitor sunitinib was identified as a common inhibitor across the three subsets. Sunitinib displayed a balanced affinity in all clusters and binds to in total 52% of all kinases with an overactivity (24 out of 46 kinases). A comparable, yet less pronounced affinity across clusters was seen for nintendanib (17 out of 46) and dasatinib (15 out of 46).

**Fig. 2.**
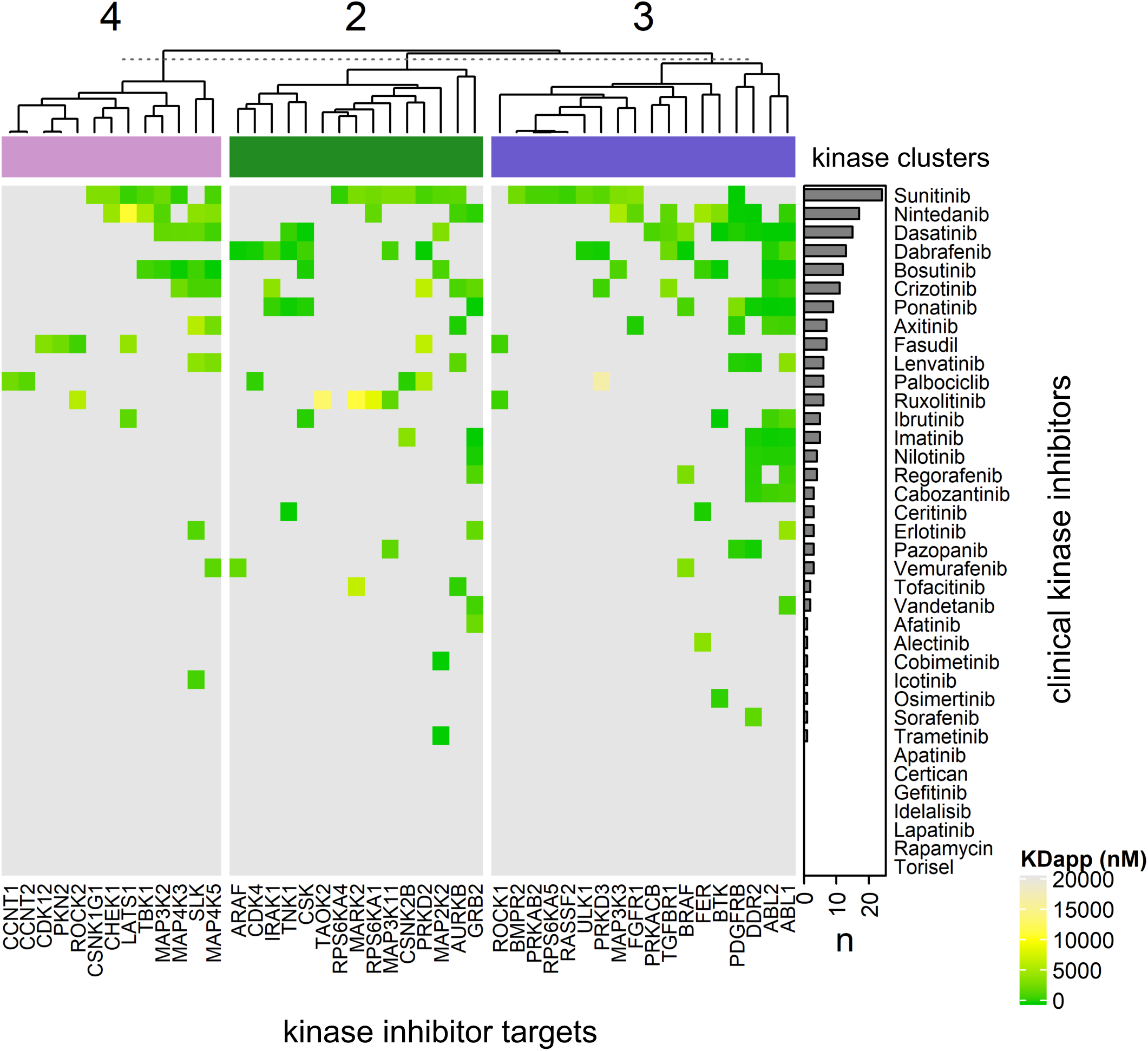
Target landscape of clinical kinase drugs in kinase-driven STADs. The heatmap includes the kinase clusters 2, 3 and 4 (n=46), corresponding to the sample cluster 2-4. The heatmap is colored according the apparent dissociation constant (KDapp) in nM. A lower value indicates a higher affinity. The row-wise barplot sums up the number of drug targets across all samples in clusters 2-4.

### Established sunitinib targets are coexpressed with cluster 3 kinases

The integrative analysis was performed based on the recently published target landscape of clinical kinases by Klaeger and colleagues. However, most efforts to understand the response to sunitinib have been based on the established and validated sunitinib targets FLT1, FLT4, KDR, PDGFRA, RET and KIT (information from drugbank.ca). Correlating the protein activity of the validated targets with the kinases belonging to clusters 2-4 showed a strong coexpression with the cluster 3 kinases (Fig. 3). This data suggests that current biomarker studies only focus on one of three kinase-driven gastric adenocarcinoma subtypes.

**Fig. 3.**
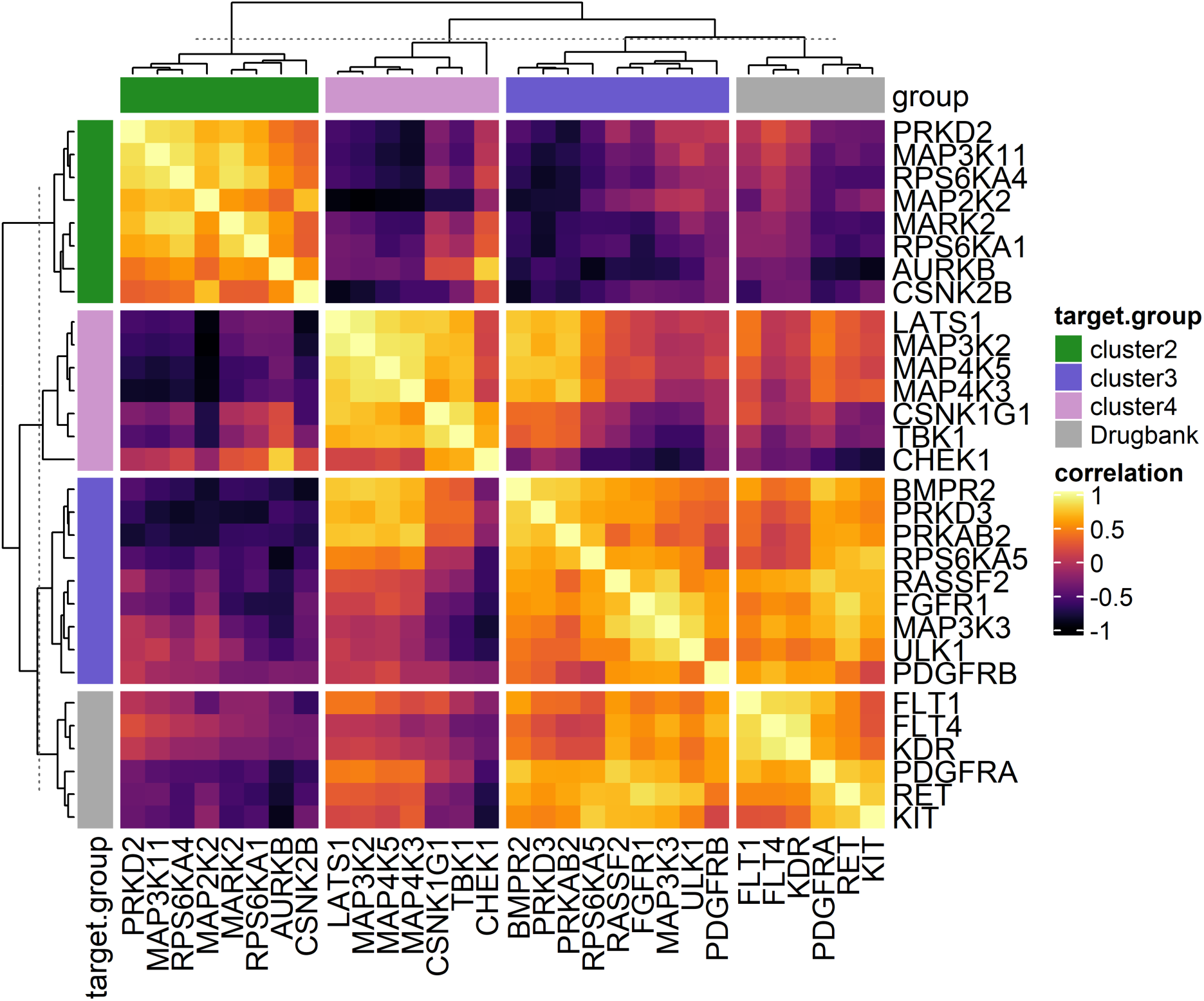
Coexpression heatmap for protein activities across sunitinib targets. In addition to the sunitinib targets revealed by the chemical proteomics screen by Klaeger and colleagues, the established and validated sunitinib targets FLT1, FLT4, KDR, PDGFRA, RET und KIT (information from drugbank.ca) are shown. The heatmap coloring corresponds to the Pearson correlation coefficient.

## Discussion

In this study, we investigated the signaling pathway dependencies in gastric adenocarcinoma in the TCGA cohort based on estimated protein activities using computational inference and integrated this with a recently published landscape of clinical kinase drugs to search for potentially effective compounds. Our analysis revealed that 43% of gastric adenocarcinomas showed elevated kinase activities and can therefore be expected to profit from kinase inhibition. The remaining 57% of samples didn’t show elevated kinase activity and should thus be more susceptible to other treatment approaches. Using hierarchical clustering those kinase-driven samples could be stratified into three groups, each group presenting mutually exclusive kinase activities. Integration with the target landscape of approved clinical kinase drugs revealed sunitinib to be a promising drug across all kinase-driven tumors.

Several studies have already tested the efficacy of sunitinib in gastric cancer in an unstratified setting. A phase II study in advanced gastric cancer employing sunitinib monotherapy as the second-line systemic treatment led to an objective response rate (ORR) of 2.6%, a stable disease (SD) rate of 32.1% and to progressive disease (PD) in 53.8% of patients with 11.5% of patients remaining unassessed (Bang et al., 2011). With 54% of patients not-benefitting from treatment, this proportion is consistent with the 57% of tumors in our analysis that were not kinase-driven. A small phase II and pharmacodynamic study in 25 patients with relapsed or refractory oesophageal or gastro-oesophageal cancers led to a SD rate of 42% and an ORR of 13% and thus provided comparable numbers (Wu et al., 2015). The authors furthermore identified higher baseline VEGFC serum levels in sunitinib responders. With sunitinib monotherapy only showing modest efficacy, sunitinib was next tested in addition to the FOLFIRI chemotherapy scheme as second- or third-line treatment for advanced adenocarcinoma of the stomach or lower esophagus (Moehler et al., 2016). Here, a similar median PFS between both groups and a trend towards prolonged median OS in the FOLFIRI/Sunitinib group was found. This study again indicates a potential outcome improvement by sunitinib that is difficult to statistically quantify in an unstratified setting. Moehler and colleagues questioned the prognostic and predictive role of serum levels of VEGFA, KDR, VEGFD and CXCL12. While serum KDR decreased after the 1^st^ cycle, the longer PFS observed at high concentrations was independent of the treatment group.

As for biomarker studies, our coexpression analysis suggest, that by assessing known sunitinib targets only approximately one third of kinase-driven tumors can be identified (corresponding to cluster 3). Hence, there is a need to also include genes from our clusters 2 and 4 in further biomarker studies. Of note, our pilot integrative kinase analysis was exclusively performed *in silico*. Further functional genomics experiments are warranted to validate the finding of kinase-driven tumors, as well as the predictive role for sunitinib response.

## Figure legends

**Fig. S1.**
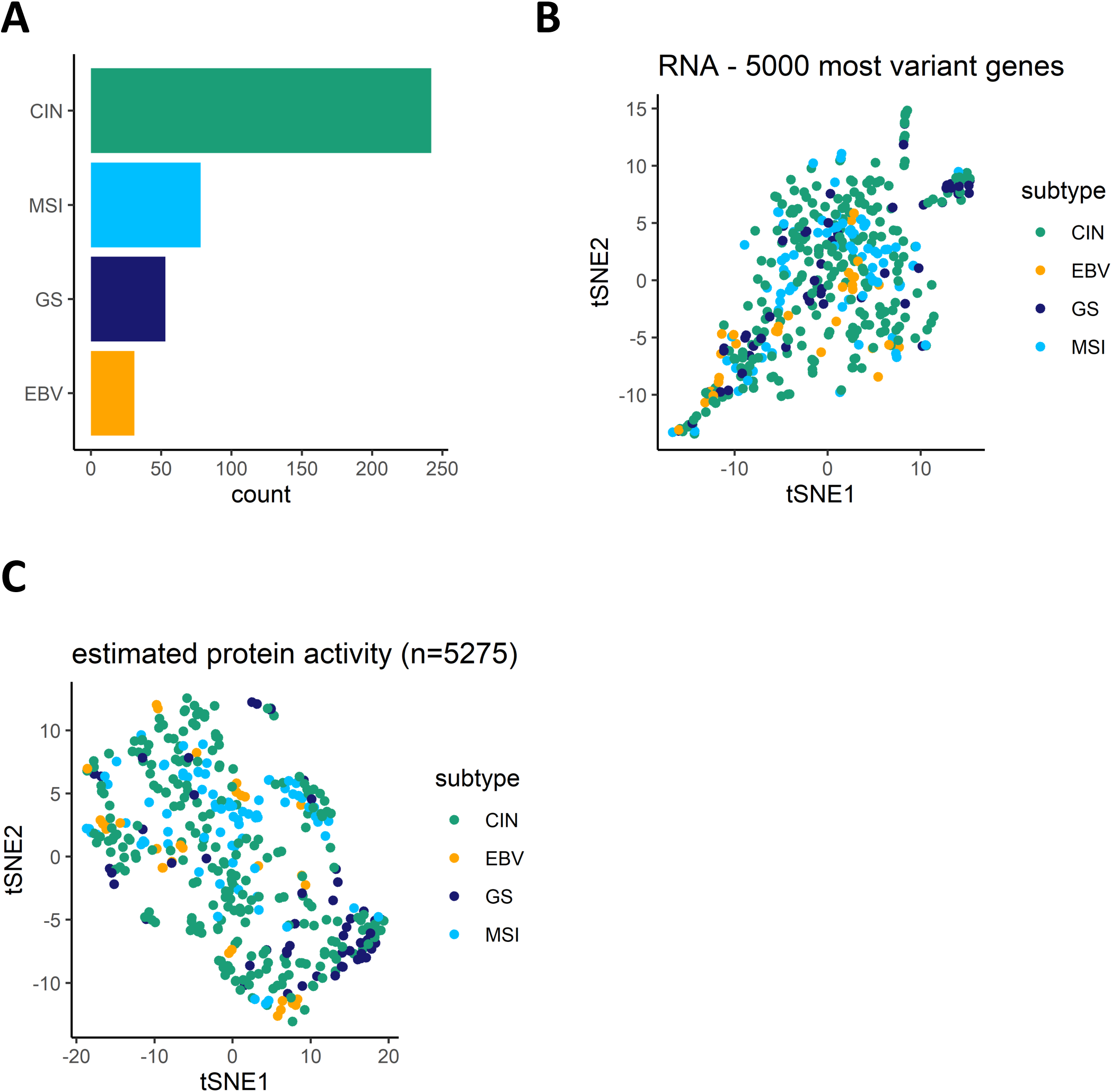
Unsupervised comparison of molecular subgroups. (A) Frequency of molecular subgroups across the TCGA cohort. (B,C) T-distributed stochastic neighbor embedding (t-SNE) plot based on (B) the 5000 most variant genes on the RNA level and (C) the 5275 genes for which an estimation of protein activity was possible. CIN, chromosomal instability; EBV, Epstein-Bar virus; GS, genome stable; MSI, microsatellite instability.

**Fig. S2.**
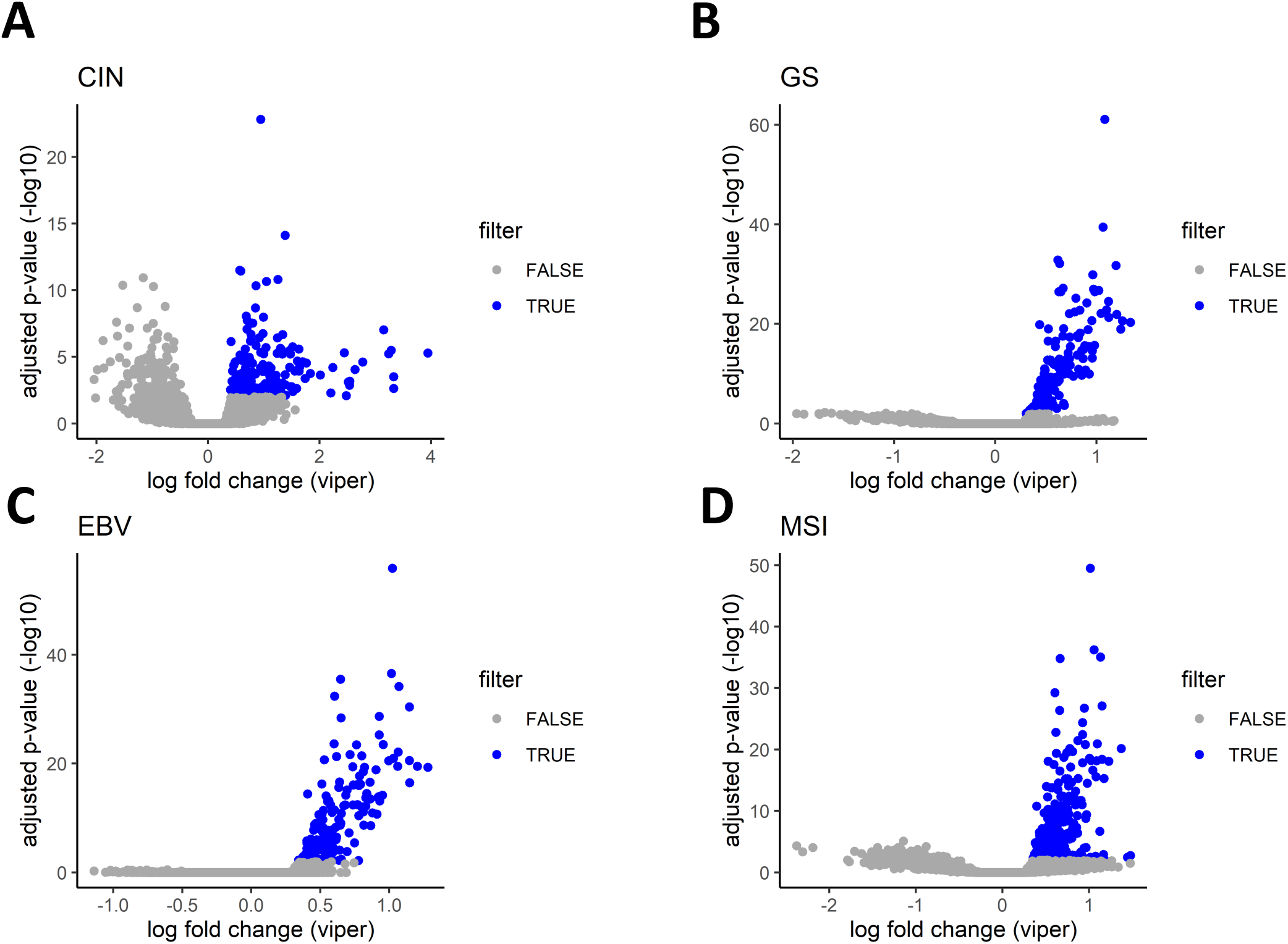
Supervised comparison of the protein activities between the molecular subgroups. **(A-D)** Volcano plots illustrating the difference between a single versus the remaining molecular subgroups. The FDR-adjusted p-value of the linear model is plotted over the log2 fold change. Points are colored according the filter (adjusted p-value <0.01 & log2 fold change > 0.25). CIN, chromosomal instability; EBV, Epstein-Bar virus; GS, genome stable; MSI, microsatellite instability.

